# Assessing the Evolutionary Trajectory of Arbuscular Mycorrhizal Conserved Genes in Seagrasses and Aquatic Close Relatives

**DOI:** 10.1101/2025.01.23.634575

**Authors:** Cassandra L. Ettinger, Jennifer Arroyo, Jason E. Stajich

**Author notes:** **Corresponding Author:** Cassandra L. Ettinger.

## Abstract

Arbuscular mycorrhizal fungi (AMF) form beneficial associations with plants, and are thought to have been critical to the adaptation of the ancestor of terrestrial plants during the transition onto land. However, the ability of AMF to associate with aquatic plants is unclear. To address this, we used 65 publicly available genomes and transcriptomes (25 freshwater, 23 terrestrial and 17 marine plants) to interrogate the genomic potential to form AMF associations in aquatic plant lineages in the order Alismatales. We explored the presence or absence of homologs of 45 genes, with a a special focus on six critical genes including three that co-evolved with AMF associations (*RAD1, STR1, STR2*) and three necessary for intracellular symbiosis (*SymRK, CCaMK/DMI3, CYCLOPS/IDP3*). Our results indicate a pattern likely consistent with independent gene losses (or extreme divergence) of symbiosis genes across aquatic lineages suggesting a possible inability to form AMF associations. However, some of these conserved genes (i.e., *CCaMK/DMI3*) are purported to function in other types of fungal symbioses, such as ectomycorrhizal symbiosis, and were observed here in a subset of aquatic lineages, including seagrasses. Overall, our findings highlight the complex evolutionary trajectories of symbiosis-related genes in aquatic plants, suggesting that while AMF associations may have been lost in certain lineages, others have genes that may allow them to form alternative fungal symbioses which may still play an underappreciated role in their ecology.

## Introduction

Arbuscular mycorrhizal fungi (AMF) are obligate mutualistic root symbionts that assist host plants in nutrient uptake in exchange for carbon compounds (Parniske, 2008; Bonfante & Anca, 2009). These fungi are thought to have been essential in facilitating the transition of ancestral plants from an aquatic to terrestrial environment (Redecker *et al*., 2000; Heckman *et al*., 2001; Wang *et al*., 2010; Humphreys *et al*., 2010). These partnerships are widespread, with the majority of flowering plants (82-85%) maintaining AMF associations, underscoring their ecological importance (Wang & Qiu, 2006; Brundrett, 2009).

AMF were historically thought not to colonize aquatic environments, but have since been observed in wetlands, estuaries, mangrove forests and freshwater ecosystems (Beck-Nielsen & Madsen, 2001; Bohrer *et al*., 2004; Šraj-Kržič *et al*., 2006; Kohout *et al*., 2012). AMF have been reported to form associations with freshwater Alismatales, an order of flowering plants that includes many aquatic lineages, although these associations appear sporadic (Clayton & Bagyaraj, 1984; Wigand & Stevenson, 1994; Khan & Belik, 1995; Beck-Nielsen & Madsen, 2001; Wang & Qiu, 2006; Šraj-Kržič *et al*., 2006; Radhika & Rodrigues, 2007; Brundrett, 2009). Notably, marine Alismatales (i.e., seagrasses) have not been observed to form AMF associations, which may be due to the additional selective pressures of the marine environment (Nielsen *et al*., 1999). Instead there have been reports of ectomycorrhizal-like fungal root symbioses in Mediterranean seagrasses, *Posidonia oceanica* and *Thalassodendron ciliatum* (Vohník et al., 2015, 2019; Borovec & Vohník, 2018; Vohník & Josefiová, 2024). Further, a recent study surveying 34 conserved plant genes involved in AMF profiled two Alismatales species, the freshwater duckweed *Spirodela polyrhiza* and the seagrass *Zostera marina*, and concluded both were likely non-mycorrhizal (NM) based on gene loss (Radhakrishnan *et al*., 2020). However, seagrasses and freshwater lineages in the Alismatales have interesting evolutionary histories which involve multiple independent transitions into aquatic ecosystems (Les *et al*., 1997; Chen *et al*., 2022). Therefore, we set out to more robustly profile the Alismatales overall to better understand how aquatic transitions might inform the evolutionary trajectory of genes involved in fungal symbiosis.

To interrogate the genomic potential to form AMF associations in aquatic plant lineages in the order Alismatales, including seagrasses and freshwater relatives, we searched for the presence or absence of key genes implicated in AMF symbiosis in a set of plant genomes and transcriptomes. We used 65 publicly available genomes and transcriptomes downloaded from NCBI GenBank (Table S1). This included 25 freshwater, 23 terrestrial and 17 marine representatives, with 49 representing members of the order Alismatales. We performed reciprocal blast searches of protein sequences to explore the presence or absence of homologs of 45 genes reported to be related to AMF symbiosis (Table S2) (Delaux *et al*., 2013, 2014; Radhakrishnan *et al*., 2020), with a special focus on six critical genes highlighted by Radhakrishnan *et al*. (2020) including three that co-evolved with AMF associations (*RAD1, STR1, STR2*) and three necessary for intracellular symbiosis (*SymRK, CCaMK/DMI3, CYCLOPS/IDP3*).

## Materials and Methods

Publicly available genomes and transcriptomes from 65 plants were downloaded from NCBI (Table S1). This included 25 freshwater, 23 terrestrial and 17 marine representatives, with 49 representing members of the order Alismatales (‘Darwin Tree of Life’; Ouyang *et al*., 2006; The International Brachypodium Initiative, 2010; Massa *et al*., 2011; Lamesch *et al*., 2011; Bennetzen *et al*., 2012; Ming *et al*., 2013, 2015; Droc *et al*., 2013; Regier *et al*., 2013; Motamayor *et al*., 2013; Wang *et al*., 2014, 2023, 2024; Schmutz *et al*., 2014; Yang *et al*., 2015; Hirsch *et al*., 2016; D’Esposito *et al*., 2016; Michael *et al*., 2017, 2021; Suzuki *et al*., 2017; Ruocco *et al*., 2017; Malandrakis *et al*., 2017; Zhang *et al*., 2017; Entrambasaguas *et al*., 2017; McCormick *et al*., 2018; Hoang *et al*., 2018, 2020; Pecrix *et al*., 2018; An *et al*., 2019; Bellinger *et al*., 2020; Yin *et al*., 2021; Ma *et al*., 2021, 2024; Park *et al*., 2021; Rowarth *et al*., 2021; Abramson *et al*., 2021; Gao *et al*., 2022; Frisse *et al*., 2022; Tang *et al*., 2022; Bayer *et al*., 2022; Ndhlovu & von der Heyden, 2022; Li *et al*., 2023; Pasaribu *et al*., 2023; Qian *et al*., 2024). We obtained the predicted AMF status for each plant family from FungalRoot v. 2 (Soudzilovskaia *et al*., 2022). FungalRoot classifies plant families as obligate arbuscular mycorrhizal (AM), non-mycorrhizal (NM), or AM-NM which refers to facultative AMs - plants that are occasionally colonized by AMF, but are often observed without AMF in other conditions. Of the 65 plants included in this work: 22 were AM, 23 were NM and 20 were AM-NM.

Using a set of 45 genes reported previously to be involved in AMF symbiosis (Table S2), including six genes, *SymR/DMI2, CCaMK/DMI3, CYCLOPS/IPD3*, GRAS transcription factor *RAD1*, and two half-ATP-binding cassette (ABC) transporters, *STR* and *STR2*, thought to be critically important for forming intracellular symbioses and AMF (Delaux *et al*., 2013, 2014; Radhakrishnan *et al*., 2020), we surveyed the genetic landscape of several plants across the order Alismatales to assess whether they have maintained the potential to form AMF associations as many species in this order have ancestors that evolutionarily transitioned from terrestrial to aquatic habitats.

We used command-line NCBI BLAST v. 2.16.0+ (Camacho *et al*., 2009) to perform reciprocal blast searches for the set of 45 genes using protein sequences from *Medicago truncatula* (v.4) as a reference, similar to searches performed in other studies of AMF (Delaux *et al*., 2014; Radhakrishnan *et al*., 2020). For annotated genomes (n=53) we searched with blastp using the protein sequences from *M. truncatula*. Top matches were then reciprocally searched back against the annotation for *M. truncatula* using blastp. While, for unannotated genomes (n=2) and transcriptomes (n=14), we instead searched with tblastn using the protein sequences from *M. truncatula*. Top matches were then reciprocally searched against the *M. truncatula* annotation using blastx. Subsequent analysis and visualization of reciprocal matches was performed in R v. 4.3.1 (R Core Team, 2021) using the tidyverse v. 2.0.0 R package (Wickham *et al*., 2019).

For the purposes of this study, we consider cases where a positive reciprocal match was made (i.e. where the initially searched *M. truncatula* gene was subsequently the top hit in the reciprocal search) to be homologous genes. Note, we did not further assess whether positive reciprocal matches represented genes that were complete or functional. Further we consider cases where an initial match was made, but the reciprocal match was negative, to potentially indicate distant homology, such as paralogs, or genes that have undergone significant evolution or divergence. Finally, we consider cases where no matches were made to most likely indicate gene loss as the ancestral state of these genes is present. Note that for unannotated genomes and transcriptomes, there are several additional reasons genes might be missed using this approach including assembly issues, transcriptional variability, abundance or long introns. However, including both unannotated genomes and transcriptomes allows us to have a more robust understanding of the Alismatales gene space using currently available public data.

## Results and Discussion

Across the 45 surveyed genes, positive reciprocal matches using protein sequences were found in 57.27 ± 3.08% of predicted AM plants, 11.78 ± 2.83% of NM-AM plants, and 18.36 ± 4.25% of NM plants (Figure 1). A similar trend emerged across habitat types: 61.26 ± 3.30% in terrestrial plants, 10.22 ± 2.97% in freshwater plants, and 14.90 ± 4.32% in marine plants. Recovery of positive reciprocal matches was highest in annotated genomes, followed by transcriptomes, and lowest in unannotated genomes (e.g., for AM plants: 82.04 ± 4.04%, 45.56 ± 4.44%, and 23.05 ± 3.23%, respectively). This suggests that efforts to expand available Alismatales assembled and annotated genomes, particularly for aquatic Alismatales, would improve detection accuracy, and likely yield additional insights into the presence and absence of these genes in aquatic lineages.

**Figure 1.**
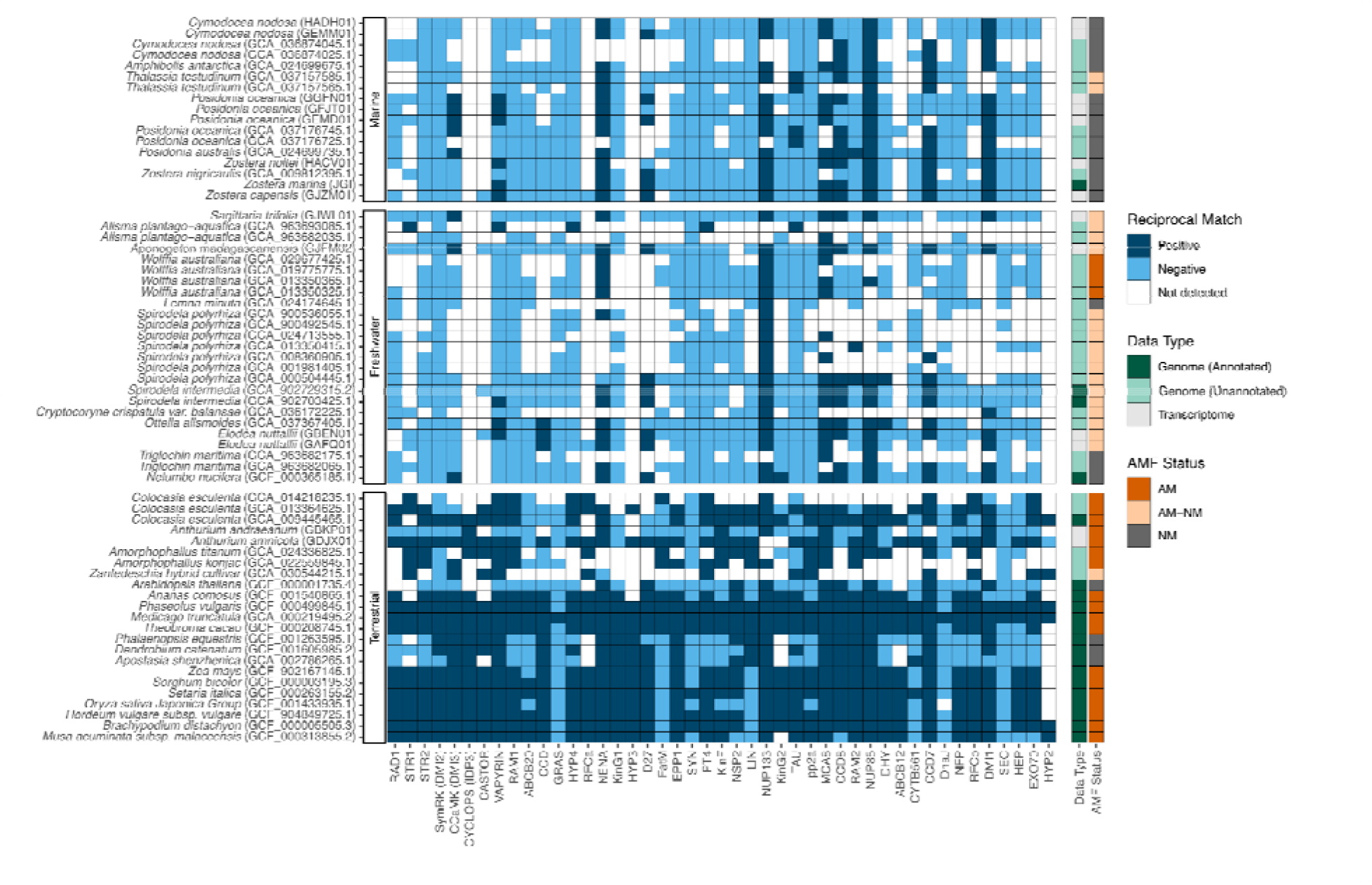
Loss of conserved symbiotic genes in aquatic plants. Heatmap showing initial blast and reciprocal blast results across marine, freshwater and terrestrial plants. Data type (genome [annotated], genome [unannotated], transcriptome) and FungalRoot v. 2 predicted AMF status (arbuscular mycorrhizal [AM], facultative arbuscular mycorrhizal [AM-NM], non-arbuscular mycorrhizal [NM]) are shown to aid interpretation.

Regardless, the lower detection of positive reciprocal matches in freshwater and marine plants suggests that these plants have lost or have experienced divergence in many of the genes involved in AMF symbiosis. These findings support the idea that plants transitioning back into aquatic environments have repeatedly lost the ability to form AMF associations, largely aligning with the NM and NM-AM predictions for these lineages from the FungalRoot database based on observational reports. Overall, our results show a pattern of likely independent loss of conserved AMF associated genes across aquatic and marine Alismatales lineages and are consistent with previous research showing convergent loss of AMF genes in other NM plants (Delaux *et al*., 2014; Radhakrishnan *et al*., 2020).

Focusing on the six key genes previously identified as vital to AMF symbiosis (Radhakrishnan *et al*., 2020), we observed similar patterns to the full gene set (Figure S1). Specifically for the three genes that co-evolved with AMF associations (*RAD1, STR1, STR2*), reciprocal matches using protein sequences were primarily found in terrestrial AM plants (83.33 ± 3.21%), with occasional matches in NM-AM freshwater plants, but none in marine plants. Radhakrishnan *et al. (2020)* suggested that the observed consistent loss of *RAD1, STR1* and *STR2* following the abandonment of AMF associations indicates there may be selection against these genes once AMF symbiosis is no longer needed. The absence of these genes in marine lineages reinforces previous observational reports that seagrasses lack AMF associations (Nielsen *et al*., 1999).

For the three genes required for intracellular symbiosis (*SymRK, CCaMK*/*DMI3, CYCLOPS/IDP3*) most reciprocal matches using protein sequences were from terrestrial plants (73.91 ± 4.35%), with only *CCaMK*/*DMI3* detected in both freshwater and marine plants. Radhakrishnan *et al*. (2020) found that these three genes were not absent from all species that have lost AMF associations and instead observed that they were maintained in species that formed other types of fungal and bacterial symbiosis. They also highlighted several additional genes (*KinF, EPP1, VAPYRIN, LIN, CASTOR*, and *SYN*) which were maintained in species capable of other types of intracellular symbioses. Of these additional genes, we only recovered positive matches to *VAPYRIN* in aquatic plants.

Our results suggest that some genes involved in AMF associations (i.e., *CCaMK/DMI3*) are retained in aquatic lineages, including seagrasses. Many of the 45 surveyed conserved genes are involved in other microbial symbioses beyond AMF (e.g., orchid mycorrhizas, ericoid mycorrhizas and legume-rhizobia symbioses) and their retention, loss or divergence may be important for understanding their interactions with other microorganisms. For example, *CCaMK/DMI3* is thought to play a role in ectomycorrhizal (ECM) symbioses (Li *et al*., 2024). In our study, we found positive reciprocal matches for *CCaMK/DMI3* in the seagrass *Posidonia oceanica*, which may relate to its reported ECM-like symbiotic relationship with the fungal root endophyte *Posidoniomyces atricolor* (Vohník et al., 2015, 2019; Borovec & Vohník, 2018). However, the function of *CCaMK/DMI3* in aquatic plants remains unclear. For example, in the aquatic *Nelumbo nucifera*, which has also been reported to have *CCaMK/DMI3*, recent work has documented a deletion in the kinase domain that likely renders it non-functional (Bravo *et al*., 2016; Radhakrishnan *et al*., 2020). These findings highlight the need for further research to confirm whether *CCaMK/DMI3* and other symbiosis-related genes detected in aquatic Alismatales remain functional and whether they have been co-opted for use in alternative symbiotic relationships.

The loss of AMF associations in plants has been speculated to be due to adaptation to nutrient-rich ecological niches, which are known to inhibit the formation of AMF (Read & Perez-Moreno, 2003). While aquatic environments are not wholly nutrient-rich, it is possible that the selective pressures of freshwater and marine life, such as low oxygen or high salinity, make it harder to maintain AMF (Nielsen *et al*., 1999; Fusconi & Mucciarelli, 2018). Additionally, many aquatic plants utilize their leaves, in addition to their roots, for uptake of nutrients which may make AMF associations, which are traditionally thought to occur in roots, less necessary (Short & McRoy, 1984; Terrados & Williams, 1997). Overall, the Alismatales are a notable group for investigating the evolution of plant-fungal symbiosis and the impact of aquatic transitions on host-microbe interactions. This work serves to highlight the need for additional genome assembly and annotation efforts for plants in this group in order to understand the molecular basis of these interactions and the functional significance of retained symbiosis-related genes.

## Acknowledgements

We would like to thank Jonathan A. Eisen for engaging in initial conversations around this work and for feedback on this manuscript. We would like to thank the Darwin Tree of Life project (https://www.darwintreeoflife.org) for making their genome assemblies publicly available without restriction.

## Competing interests

The authors declare no competing interests.

## Author contributions

CLE conceived and designed the analysis, analyzed the data, prepared figures and/or tables, wrote and reviewed drafts of the paper. JA prepared figures and/or tables, and wrote and reviewed drafts of the paper. JES reviewed drafts of the paper.

## Data availability

All code used in this work has been deposited on Github (casett/Seagrass_AMF) and archived in Zenodo (DOI: 10.5281/zenodo.14218123). The genomic and transcriptomic data used in this work are publicly available and were obtained from NCBI GenBank (Table S1).

## Funding sources

CLE was supported by the National Science Foundation (NSF) under a NSF Ocean Sciences Postdoctoral Fellowship (Award No. 2205744). JA was supported by the NSF REU The National Summer Undergraduate Research Project, Award No. 2149582. JES is a CIFAR Fellow in the program Fungal Kingdom: Threats and Opportunities and was partially supported by NSF awards EF-2125066 and IOS-2134912. Computations were performed using the computer clusters and data storage resources of the UC Riverside HPCC, which were funded by grants from NSF (MRI-2215705, MRI-1429826) and NIH (1S10OD016290-01A1). The funders had no role in study design, data collection and analysis, decision to publish, or preparation of the manuscript.

## Notes

### Competing Interest Statement

The authors have declared no competing interest.

### Summary of Updates

Corresponding author email updated; no manuscript changes

