## Supplemental Figures and Table Legends for "Assessing the Evolutionary Trajectory of Arbuscular Mycorrhizal Conserved Genes in Seagrasses and Aquatic Close Relatives"

^#^**Corresponding Author:**

Cassandra L. Ettinger

**Supplemental Table Legends and Figures:**

**Table S1**. Genome and transcriptome information. Here we provide details on each dataset obtained from NCBI including plant species, order, family, BioProject number, and assembly accession number. We also report the habitat of each species (marine, freshwater or terrestrial), FungalRoot v. 2 AMF predicted status, annotation status and appropriate data citation.

**Table S2.** Conserved symbiosis genes. The table reports the gene IDs of all genes used in this work, the respective gene code in the *M. truncatula* v. 4 reference genome and the protein sequence that was used for all blast searches in this work.

**Figure S1.** Loss of six critical conserved symbiotic genes in aquatic plants. Heatmap showing initial blast and reciprocal blast results across marine, freshwater and terrestrial plants for six genes including three thought to have co-evolved with AMF associations (*RAD1, STR1, STR2*) and three necessary for intracellular symbiosis (*SymRK, CCaMK, CYCLOPS*). Data type (genome [annotated], genome [unannotated], transcriptome) and FungalRoot v.2 predicted AMF status (arbuscular mycorrhizal [AM], facultative arbuscular mycorrhizal [AM-NM], non-arbuscular mycorrhizal [NM]) are shown to aid interpretation.

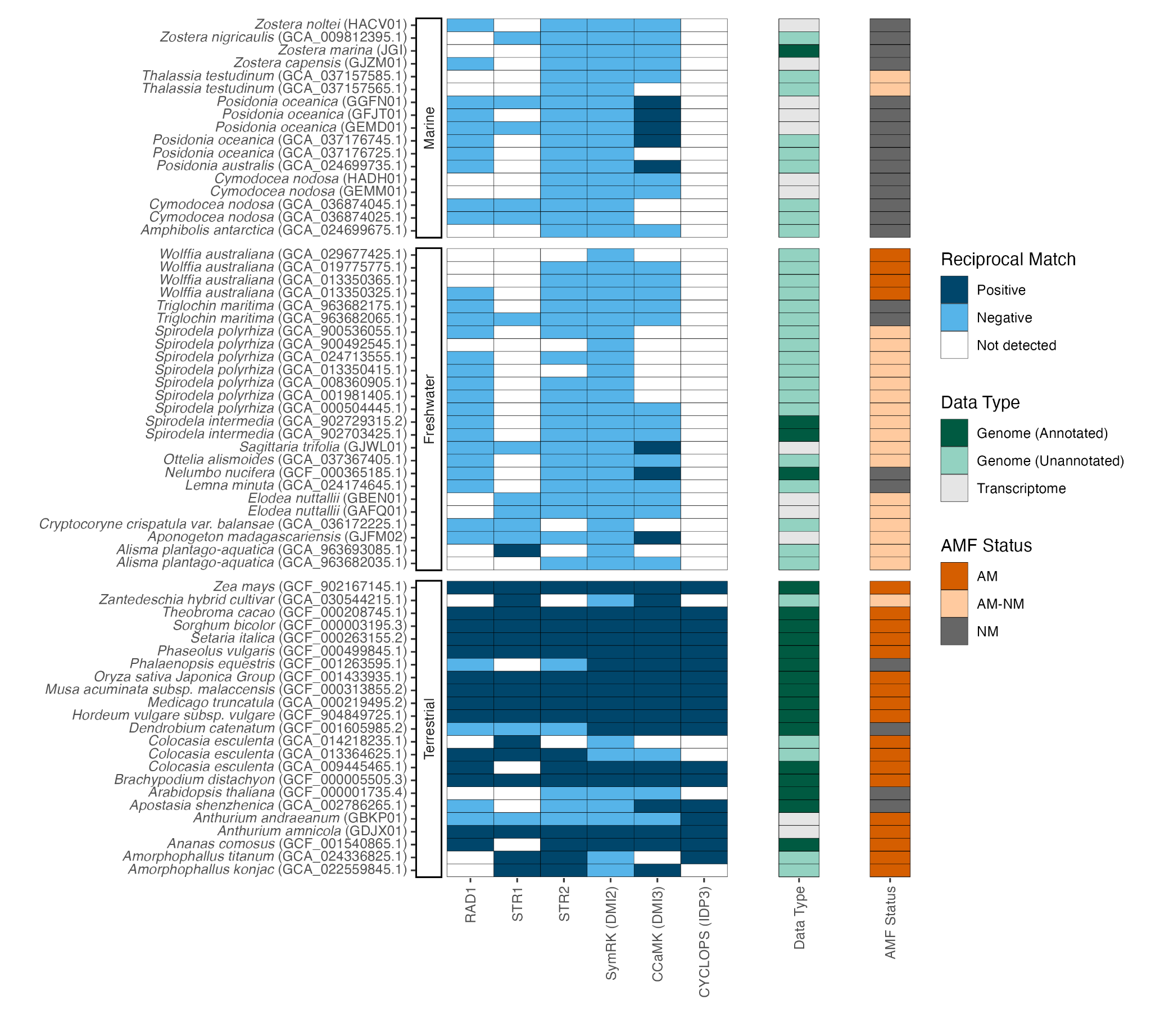
